# *Xanthomonas citri* pv. *lagerstroemium*, description of a new pathovar causing leaf spot on crape myrtle

**DOI:** 10.1101/2025.02.28.640647

**Authors:** Izabela M. Duin, David Ritchie, Evan Braswell, Jennie Fagen

## Abstract

Bacterial leaf spot caused by *Xanthomonas* was reported in 2014 as a new disease of crape myrtle. Unfortunately, this foundational strain was lost, preventing further experimentation, sequencing of the genome, and phylogenetic analysis. This work describes a collection of *Xanthomonas* strains isolated from angular leaf spot lesions on crape myrtle in North Carolina from 2014 to 2023. This study includes full reference genomes, as well as re-fulfillment of Koch’s postulates. Genomes were obtained with hybrid whole genome sequencing using Illumina and Nanopore and assembled to develop robust genomic resources for these disease-causing strains. The completed genomes support inclusion of the strains in the *X. citri* species group; however, both phylogenetic analysis and the identification of a novel plant host suggest the creation of the new pathovar *Xanthomonas citri* pv. *lagerstroemium*.

## Short Communication

Crape myrtle (*Lagerstroemia indica* and hybrids) is a globally relevant ornamental woody plant and is the representative species of the genus *Lagerstroemia*. In 2019, the United States horticultural industry produced more than 3 million trees with a value exceeding US $69 million; over 4 million dollars of which were retail sales (USDA NASS 2020). Crape myrtles are popular with homeowners and commercial landscapers due to their long-lasting abundant blooms throughout the hot summer months, as well as historically low numbers of significant pests and diseases (Dirr 2009). Scale insect, *Acanthococcus lagerstroemiae*, were first found in crape myrtle in Texas in 2004 (Merchant et al. 2004) and has since spread across twelve states in the Southeast US (Wang et al. 2016). Powdery mildew and *Cercospora* leaf spot are the primary reported foliar diseases for crape myrtle (Chappell et al. 2012). These pests and pathogens both detract from the visual appeal of these ornamental trees and, in severe cases, can lead to weakening and decline over multiple seasons (Chappell et al. 2012).

A *Xanthomonas species* causing bacterial leaf spot on crape myrtle was reported by Babu et al. (2014). The disease was observed in numerous nurseries in Florida in 2011 and field surveys at the University of Florida, North Florida Research and Education Center, Quincy, FL in 2012 and 2013. The symptoms were characterized as dark brown, angular to irregularly shaped, oily-looking spots surrounded by yellow halos. Symptoms primarily occurred on mature leaves and in severe cases rapid defoliation of susceptible cultivars, such as ‘Arapaho’ and ‘Tuscarora’ has been observed (Babu et al 2014). Isolates from crape myrtle exhibiting angular leaf spot symptoms were collected from 2014 - 2023 primarily from nursery stocks at garden centers including plants known to have originated in the Florida panhandle where the pathogen was first described (Table 1). Additionally, established symptomatic trees were identified at the North Carolina Sandhills Research Station near Jackson Springs, NC (Table 1). For all symptomatic plants, a section of leaf tissue was excised using a sterile scalpel, macerated in sterile water, and streaked across sucrose peptone agar (SPA). After 72 h incubation at 28°C, the plates were dominated by yellow mucoid colonies consistent with xanthomonads grown on SPA. Single colonies were subsequently isolated and stored at -80ºC in 20% glycerol.

**Table 1.**
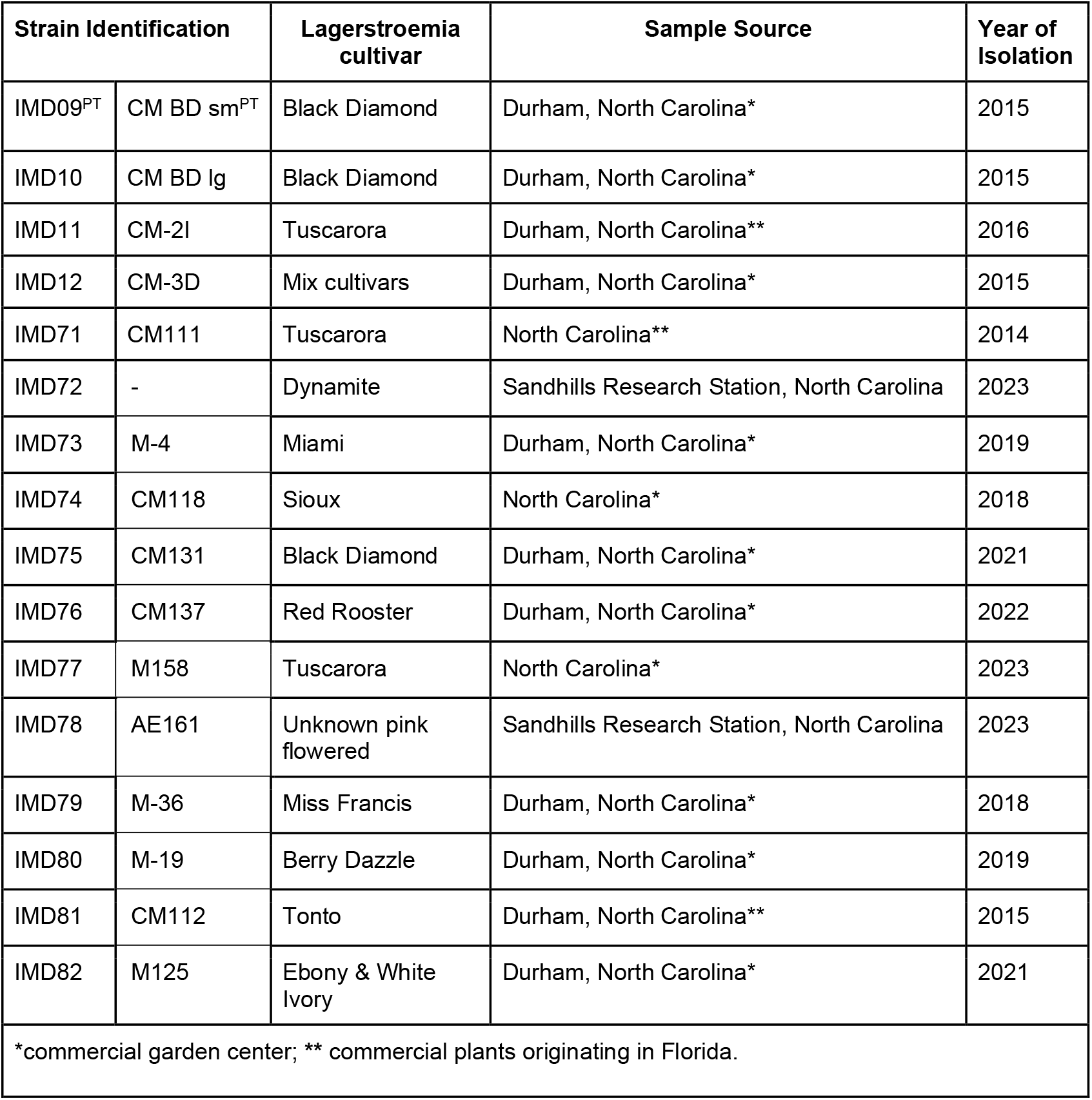
Collection location and year for isolates from *Xanthomonas*-symptomatic crape myrtle.

| Strain Identification |  | Lagerstroemia cultivar | Sample Source | Year of Isolation |
| --- | --- | --- | --- | --- |
| IMD09<sup>PT</sup> | CM BD sm<sup>PT</sup> | Black Diamond | Durham, North Carolina* | 2015 |
| IMD10 | CM BD lg | Black Diamond | Durham, North Carolina* | 2015 |
| IMD11 | CM-2I | Tuscarora | Durham, North Carolina** | 2016 |
| IMD12 | CM-3D | Mix cultivars | Durham, North Carolina* | 2015 |
| IMD71 | CM111 | Tuscarora | North Carolina** | 2014 |
| IMD72 | - | Dynamite | Sandhills Research Station, North Carolina | 2023 |
| IMD73 | M-4 | Miami | Durham, North Carolina* | 2019 |
| IMD74 | CM118 | Sioux | North Carolina* | 2018 |
| IMD75 | CM131 | Black Diamond | Durham, North Carolina* | 2021 |
| IMD76 | CM137 | Red Rooster | Durham, North Carolina* | 2022 |
| IMD77 | M158 | Tuscarora | North Carolina* | 2023 |
| IMD78 | AE161 | Unknown pink flowered | Sandhills Research Station, North Carolina | 2023 |
| IMD79 | M-36 | Miss Francis | Durham, North Carolina* | 2018 |
| IMD80 | M-19 | Berry Dazzle | Durham, North Carolina* | 2019 |
| IMD81 | CM112 | Tonto | Durham, North Carolina** | 2015 |
| IMD82 | M125 | Ebony & White Ivory | Durham, North Carolina* | 2021 |
| *commercial garden center; ** commercial plants originating in Florida. |  |  |  |  |

Total bacterial DNA was isolated for the 16 isolates from overnight bacterial cultures using the Quick-DNA Fungal/Bacterial Miniprep Kit (Zymo Research Corp, Irvine, CA, USA) following the manufacturer’s instructions and quantified using Qubit (ThermoFisher Scientific, Waltham, MA, USA). Preliminary identification via 16S rRNA gene sequence was conducted for four isolates (CM BD sm, CM BD lg, CM-2I and CM-3D) using primers 8F (Edwards et al. 1989) and 1492R (Stackebrandt and Liesack 1993). BLASTn was used to query the non-redundant nucleotide NCBI database (Altschul et al. 1990) and the sequences were found to share 99-100% ID with *Xanthomonas citri* subsp. *citri* strain Xcc49 and *Xanthomonas citri* strain CJHCd004. Differentiation of *Xanthomonas citri* at the subspecies and pathovar level necessitated additional genome-level analysis.

Unfortunately, the strains first described by Babu et al (2014) were lost in 2018 due to hurricane damage. Although the strains originally described by Babu et al. 2014 were not fully sequenced, 16S rRNA gene and partial 16S-23S internal transcribed spacer regions were available for comparison (Accessions KF926678, KF926679, KF926680, KF926681, and KF926682). These sequences were compared to strain CM BD sm using BLASTn (Altschul et al. 1997). The shared sequence identities were 2397/2398bp (99%), 2397/2403bp (99%), 2404/2408bp (99%), 2405/2406bp (99%) and 2410/2412bp (99%) respectively.

Both short and long read shotgun sequencing libraries were prepared for all 16 isolates at the Microbial Genome Sequencing Center (Pittsburgh, PA, USA); using the Illumina DNA Prep kit (Illumina, Inc., San Diego, CA, USA), IDT 10bp UDI indices and the Oxford Nanopore Technologies (ONT) Ligation Sequencing Kit (SQK-NBD114.24) with NEBNext® Companion Module (E7180L), respectively. The samples were sequenced on the Illumina NextSeq 2000 platform (2 × 151 bp) and Nanopore R10.4.1 flow cells and MinION Mk1B device. Post-sequencing, Guppy (v6.3.8) (Oxford Nanopore Technologies, Oxford, UK) was used for basecalling (SUP) and demultiplexing for ONT. Quality control and adapter trimming was performed with bcl-convert (Illumina) and porechop (Bonenfant et al. 2023) for Illumina and ONT sequencing, respectively.

The complete genome of strain CM BD sm was assembled, rotated, trimmed, and circularized with the Trycycler v.0.5.3 (Wick et al. 2021) long-read consensus assembler using default parameters. Briefly, long reads were assembled with Flye, miniasm and minipolish, and Raven (Kolmogorov et al. 2019; Wick and Holt 2019; Vaser and Šikić 2021). The resulting assembly was polished with Illumina reads using polypolish (Wick and Holt 2022) and POLCA (Zimin and Salzberg 2020). The final assembly of the strain CM BD sm yielded a circular, 5,053,938-bp chromosome (∼250× coverage) with a GC content of 64.7 (Table 2). The assembly had a 98.75% average nucleotide identity (ANI) (Yoon et al. 2017) to the *X. citri* subsp. *citri* MN12 reference genome ASM96121v1 (Figure 1). Notably, no extrachromosomal genomic elements were detected in strain CM BD sm.

**Table 2.** Description of the genome sequence of *Xanthomonas citri* pv. *lagerstroemium* isolated from crape myrtle leaves.

|  | **CM BD sm<sup>PT</sup>** | **CM BD lg** | **CM-2I** | **CM-3D** | **CM111** | **IMD72** | **M-4** | **CM118** |
| --- | --- | --- | --- | --- | --- | --- | --- | --- |
| Size (bp) | 5,053,938 | 5,010,211 | 5,134,735 | 5,136,802 | 5,013,743 | 5,041,164 | 5,042,630 | 5,290,758 |
| GC content | 64.7 | 64.8 | 64.8 | 64.8 | 64.8 | 64.7 | 64.7 | 64.6 |
| Coverage (x) | 250 | 206 | 197 | 114 | 4,257 | 5,497 | 8,906 | 2,099 |
| Number of contigs | 1 | 50 | 51 | 45 | 44 | 43 | 47 | 58 |
| Number of subsystems | 326 | 326 | 327 | 329 | 326 | 326 | 326 | 329 |
| Number of Coding Sequences | 4,603 | 4,577 | 4,723 | 4,731 | 4,594 | 4,641 | 4,622 | 4,952 |
| Number of RNAs | 61 | 56 | 56 | 56 | 56 | 56 | 56 | 53 |
| GenBank accession number | CP114795 | JBGLNA000000000 | JBGLMZ000000000 | JBGLMY000000000 | JBGLMX000000000 | JBGLMW000000000 | JBGLMV000000000 | JBGLMU000000000 |
|  | **CM131** | **CM137** | **M158** | **AE161** | **M-36** | **M-19** | **CM112** | **M125** |
| Size (bp) | 5,082,859 | 5,175,964 | 5,056,978 | 5,200,315 | 5,202,621 | 5,255,613 | 5,284,769 | 5,070,018 |
| GC content | 64.7 | 64.6 | 64.7 | 64.6 | 64.7 | 64.5 | 64.4 | 64.7 |
| Coverage (x) | 4,924 | 4,650 | 3,982 | 3,334 | 3,256 | 2,480 | 2,379 | 2,918 |
| Number of contigs | 47 | 50 | 43 | 53 | 44 | 55 | 60 | 45 |
| Number of subsystems | 326 | 326 | 326 | 327 | 329 | 326 | 329 | 326 |
| Number of Coding Sequences | 4,676 | 4,807 | 4,639 | 4,799 | 4,809 | 4,915 | 4,877 | 4,655 |
| Number of RNAs | 56 | 56 | 56 | 55 | 56 | 55 | 78 | 56 |
| GenBank accession number | JBGLMT000000000 | JBGLMS000000000 | JBGLMR000000000 | JBHJCZ000000000 | JBGLMQ000000000 | JBGLMP000000000 | JBHJCY000000000 | JBGLMO000000000 |

**Figure 1.**
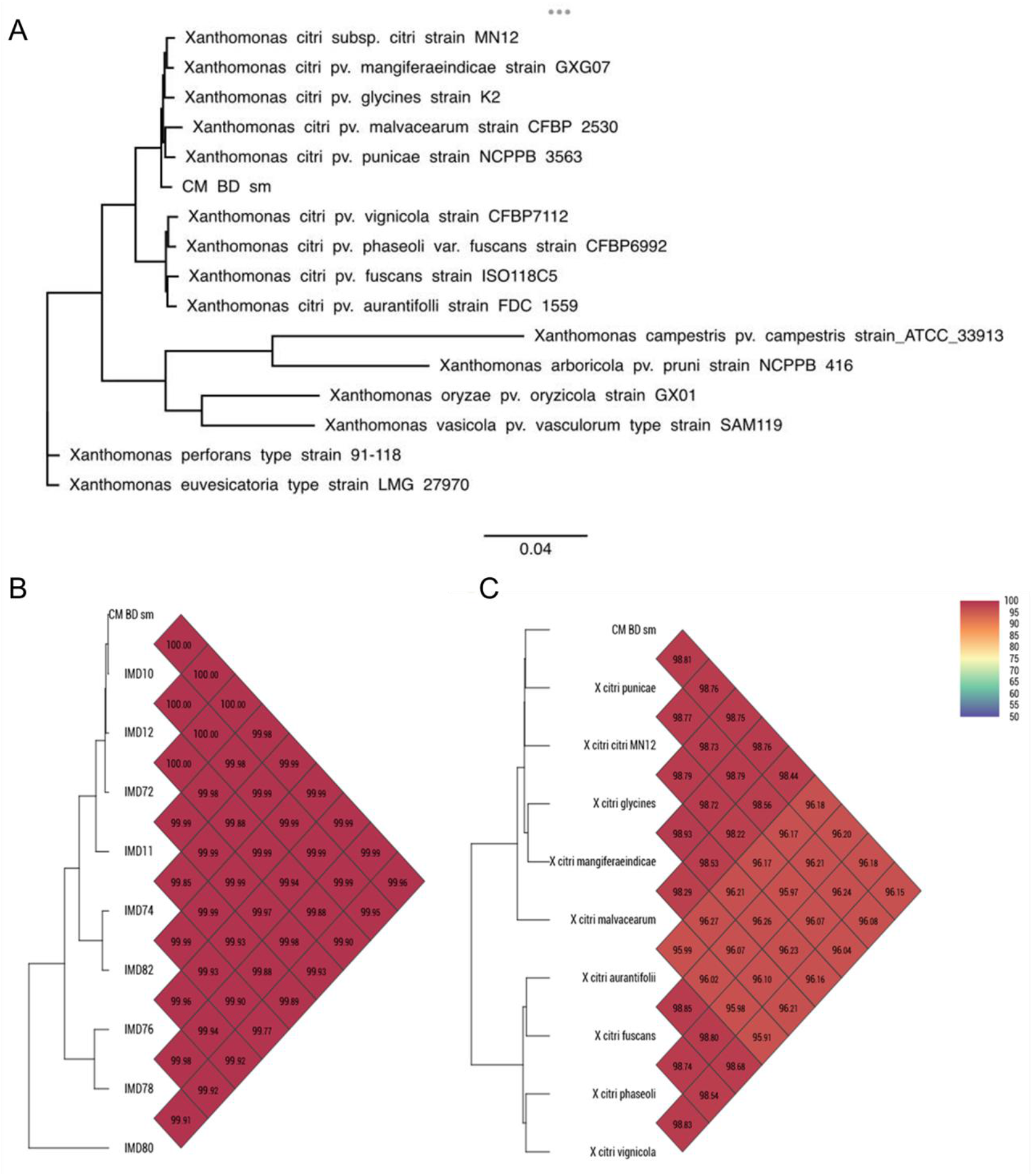
Relationships among *Xanthomonas citri* pv. *lagerstroemium* isolates and *Xanthomonas* strains. A. Maximum likelihood phylogenetic tree inferred for 16 *Xanthomonas* strains using the core genome alignment, including *Xanthomonas citri* pv. *lagerstroemium* strain CM BD sm isolated from crape myrtle. B. Average Nucleotide Identity (ANI) of ten representative *Xanthomonas citri* pv. *lagerstroemium* isolated from crape myrtle used in this study. C. ANI of representative strain *Xanthomonas citri* pv. *lagerstroemium* (CM BD sm) isolated from crape myrtle with *Xanthomonas citri* from different pathovars.

Partial genome sequences were generated for an additional 15 isolates on the Illumina NextSeq 2000 platform, as described above. The draft genomes were assembled using CLC Workbench Software. Each genome was between 5.0 to 5.2 Mb, with GC contents of 64.7 (+/-0.3) % (Table 2). All genomes were annotated using the RAST workflow (Overbeek et al. 2014). Default parameters were used for all software.

Publicly available whole genome sequences for 15 disparate *Xanthomonas* strains were obtained from NCBI GenBank. Genome annotations were performed using prokka v. 1.14.6 with default parameters to generate gff files for each strain (Seemann 2014). A pangenome was constructed for the strains using the program Roary (Version 3.13.0) (Page et al., 2015). The core genome alignment was then used to infer a maximum likelihood phylogeny using iqtree v. 2.3.4 first to determine the best fitting nucleotide substitution model based on Bayesian information criterion scores, then repeated using the selected model with bootstrapping and the approximate likelihood ratio test set to 1000 (Nguyen et al. 2015) (Figure 1A). Additionally, OrthoANI (Lee at al. 2016) was used to compare the average nucleotide identity (ANI) of the novel isolates from Crapemyrtle and the *Xanthomonas citri* group. All 16 isolates shared more than 99.87% ANI among themselves and more than 96% with other strains from the *Xanthomonas citri* species complex (Figure 1B, 1C). As *Lagerstroemia* had not been previously established as a host of *Xanthomonas citri*, we suggest the creation of pathovar, *X. citri* pv. *lagerstroemium* (Xclag), of which strain CM BD sm is the representative.

Pathogenicity tests were performed on potted crape myrtle cv. Tuscarora using three representative bacterial strains: CM BD sm, CM111, and IMD72. Bacteria were grown on nutrient agar plates for 2 days and a 10^8^ CFU/ml suspension in sterile deionized water was sprayed on the foliage to drip. The plants were covered with transparent plastic bags for 24 h before and 48 h after inoculation. Control plants were sprayed with sterile water only. The inoculated plants were maintained in a greenhouse at 28-34°C for 21 days. Plants did not receive supplemental nutrition, and all treatments developed minor interveinal chlorosis.

Symptoms of water soaked, dark brown, angular to irregularly shaped lesions were observed only on all the inoculated plants after 11 days (Figure 2). At 14 days after inoculation, the size of the lesions increased, and abscission was observed in the most heavily infested leaves. Symptoms at 21 days were consistent with the largest lesions observed at the time of the initial isolation and chlorotic halos were observed as described by Babu et al. 2014 (Figure 2). The bacterium was re-isolated from the inoculated symptomatic plants as described above and Illumina genome sequencing was performed, as described above, resulting in 99.90% of similarity with the bacteria strain inoculated, thus fulfilling Koch’s postulates.

**Figure 2.**
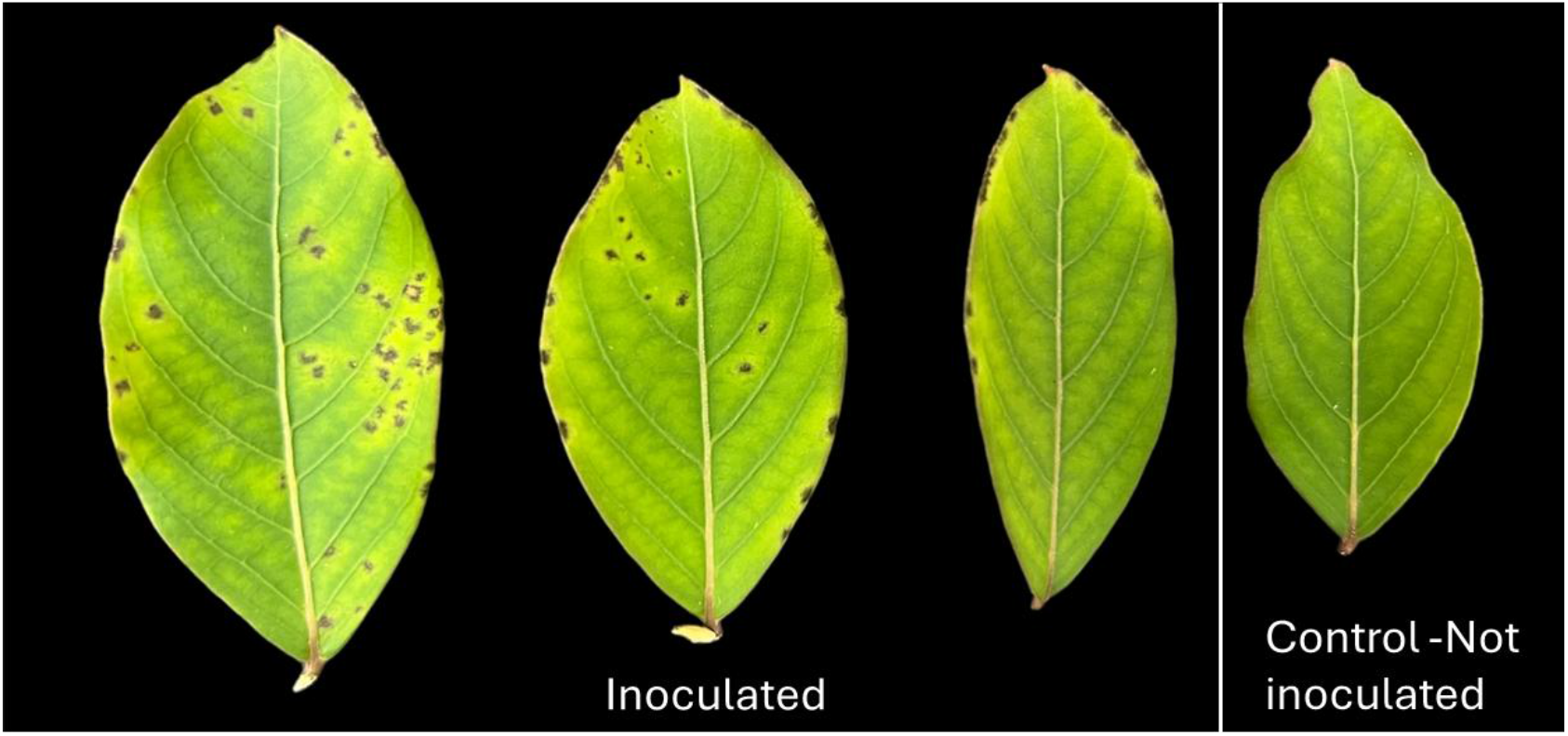
Photos of symptoms on crape myrtle cv. Tuscarora caused by CM BD sm^PT^ 22 days after inoculation. From left to right, three inoculated leaves with different levels of disease severity and one control, not inoculated leaf with no symptoms.

The new pathovar *Xanthomonas citri* pv. *lagerstroemium* expands the known host range of *Xanthomonas citri* strains. Furthermore, the newly available genomes serve as a valuable resource for evolutionary studies and facilitate investigations into the genomic factors influencing host range and virulence, thereby enhancing our understanding of pathogen adaptation and disease progression (Ochman et al. 2000; Medini et al. 2005) Enhanced awareness of this pathogen is crucial for developing disease-resistant plant cultivars and minimizing unnecessary fungicide applications, thereby reducing economic losses and environmental impact in both nursery production and landscape management (McGovern 2015; Bebber et al. 2019).

## Data availability

Sequences of the complete genome of the strain IMD09 have been submitted to GenBank under the following accession numbers: genome, CP114795; raw reads, SRR22777638 and SRR22777639, for Illumina and Nanopore, respectively; BioProject, PRJNA913162; and BioSample, SAMN32277936. The GenBank accession numbers for the draft genomes can be found in Table 2, BioProject, PRJNA1147685 and BioSample, SUB14654168.

